# EditFinder: A Sequence-Based Model for A-to-I RNA Editing Identification and Characterization of Genetic Variant Effects

**DOI:** 10.64898/2026.11.23.768754

**Authors:** Yan Zeng, Qinfeng Liu, Guisheng Liao, Xiaobo Zhou, Liyu Huang, Sijia Wu

## Abstract

A-to-I RNA editing is a widespread post-transcriptional modification that contributes to transcriptomic diversity and has been implicated in various diseases. However, incomplete reference resources for RNA editing and limited research into its genetic regulation hinder investigations into RNA editing-mediated disease mechanisms. To address these limitations, EditFinder, a sequence-based model for identifying A-to-I RNA editing sites and assessing the effects of genetic variants on RNA editing, is developed. EditFinder demonstrates superior performance on both an independent reference validation dataset and a read-based sequencing dataset. Its genome-wide application generates an expanded resource of candidate A-to-I RNA editing sites, providing a foundation for comprehensive exploration and functional annotation of RNA editing. Beyond site identification, EditFinder enables systematic assessment of the position- and nucleotide-dependent effects of genetic variations on predicted editing probabilities, highlighting the particularly strong influence of the 3’-adjacent nucleotide. Analysis of REDIportal-documented editing sites further suggests that genetic variations in flanking sequences may alter the predicted editing status of approximately 10% of the entries. Moreover, changes in EditFinder-predicted editing probabilities may reflect trends in SNP-associated variation in RNA editing frequency. Collectively, EditFinder provides a valuable resource for investigating RNA editing and its genetic regulation related to human diseases.

## 1. Introduction

A-to-I RNA editing is a widespread post-transcriptional modification that expands molecular diversity [1; 2] and has been implicated in complex diseases [3; 4]. Accurate identification of RNA editing sites ideally requires both whole-genome sequencing (WGS) and RNA-sequencing (RNA-seq) data from the same samples, or, when WGS data are unavailable, RNA-seq alone combined with stringent filtering strategies [5; 6]. The matched WGS and RNA-seq data are rarely available in most studies, limiting the applicability of the former approach. RNA-seq-based approaches commonly rely on previously catalogued editing sites, such as those in REDIportal [7], and remove variants overlapping genetic variation databases such as dbSNP to minimize potential interference from genomic SNPs. However, these resources are inherently dependent on the available sequencing datasets and reference annotations and therefore cannot provide a comprehensive representation of A-to-I RNA editing across genomic sequence contexts. Such stringent filtering may also exclude genuine disease-associated or individual-specific editing events, thereby limiting the discovery of previously uncharacterized RNA editing sites. Thus, a comprehensive and reference-genome-aware RNA editing resource is needed to systematically characterize A-to-I RNA editing.

Beyond accurate identification of RNA editing sites, understanding how genomic variation regulates RNA editing is important for elucidating the genetic determinants of post-transcriptional regulation and may provide insights into disease mechanisms and therapeutic responses. Previous studies have investigated associations between SNPs and RNA editing using statistical approaches, including editing quantitative trait locus (edQTL) analyses [8–10]. However, these association-based approaches are constrained by the quality, coverage, and allele frequencies of both SNP and RNA editing measurements. To obtain robust associations, RNA editing events and genetic variants are typically subjected to multiple quality-control and informativeness criteria, including missingness, editing-frequency variation, genotype quality, sequencing depth, minor allele frequency, and Hardy–Weinberg equilibrium, which can substantially reduce the number of variants and editing events available for association analysis. These stringent filtering procedures limit the ability to comprehensively characterize the effects of genetic variation on RNA editing. In addition, the effects of multiple SNPs on individual RNA editing sites are difficult to evaluate using conventional association-based approaches. Moreover, statistical associations alone provide limited information about the sequence-level mechanisms through which individual SNPs may alter the recognition or editing of neighboring adenosines. Therefore, a systematic approach is needed to comprehensively characterize the effects of genomic variants on RNA editing site across local sequence contexts.

To address these challenges, a model was first established using genomic reference sequences to distinguish SNP-derived nucleotide variations from A-to-I RNA editing, enabling genome-wide identification of A-to-I RNA editing sites. This model was further used to systematically evaluate how specific SNPs at upstream and downstream positions within the local sequence context affect the probability of A-to-I RNA editing. This framework provides a comprehensive reference resource for A-to-I RNA editing and enables systematic characterization of local genomic variation effects on RNA editing across the genome.

## 2. Results

### 2.1. EditFinder demonstrates superior performance in A-to-I RNA editing detection

A major challenge in A-to-I RNA editing detection is distinguishing genuine RNA editing events from nucleotide changes arising from genomic SNPs. To address this challenge, EditFinder was established as a sequence-based model that uses reference genome sequences surrounding RNA editing sites and SNPs as input. The model was trained using stringently filtered A-to-I RNA editing sites as positive samples and genomic A-to-G SNPs as negative samples. Filtering procedures were designed to minimize overlap between the positive and negative datasets and to reduce the confounding effects of local variations on sequence features in both classes (Figure 1A). Chromosome-level separation of the development and test sets was adopted to evaluate model performance on independent genomic sequences (Table S1), with the development set subjected to ten-fold cross-validation for model training. The overall architecture of EditFinder integrates sequence-based convolutional features with sequence similarity information for A-to-I RNA editing prediction (Figure 1B).

**Figure 1.**
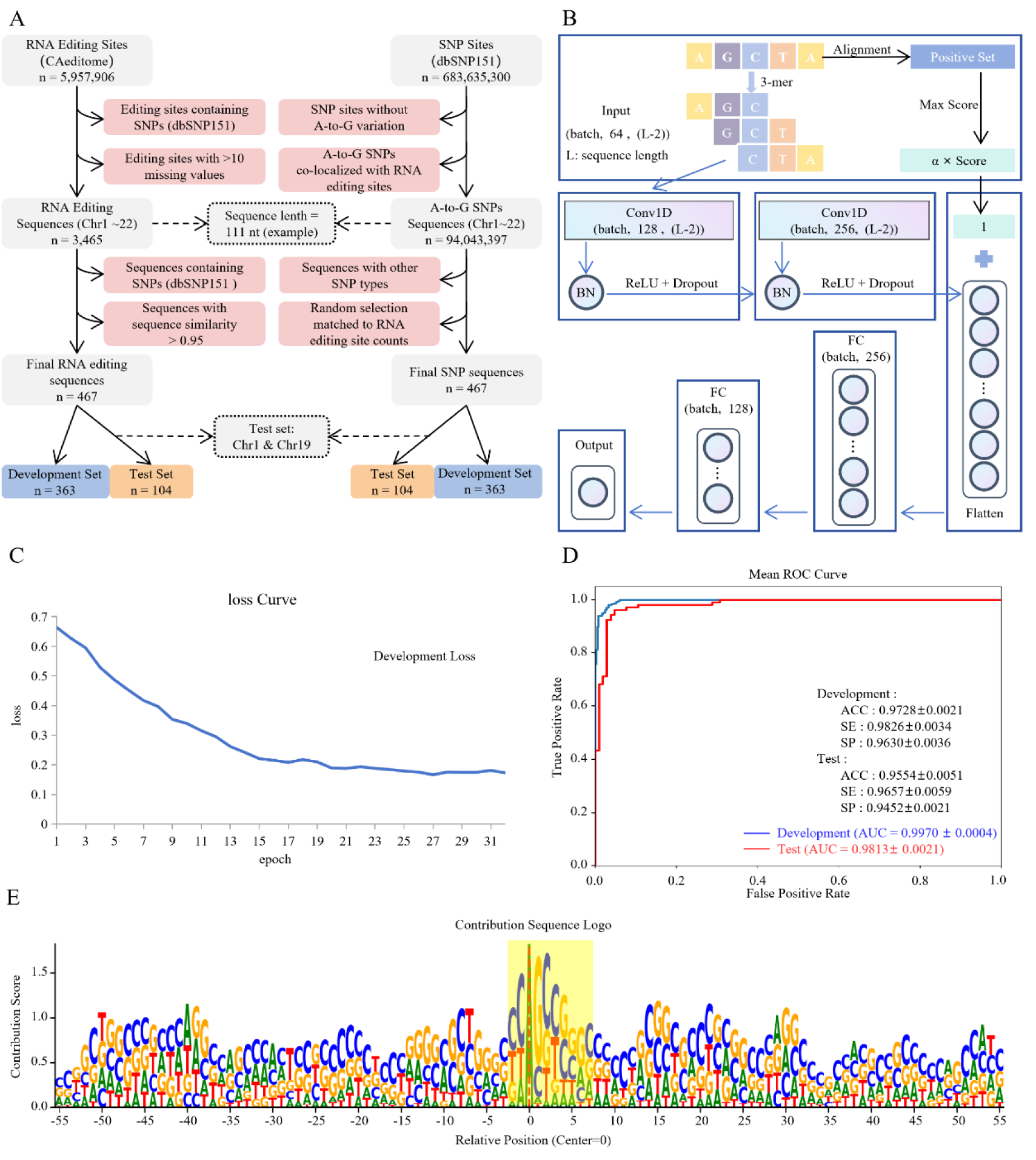
EditFinder model. (A) Data preparation pipeline for positive (RNA editing) and negative (SNP) samples. (B) Architecture of EditFinder. (C) Training process for the model. (D) Performance of the model. (E) Motif enrichment for RNA editing identification using DeepLIFT. The height of each nucleotide is proportional to its contribution score at that position. ROC: receiver operating characteristic curve, ACC: accuracy, SE: sensitivity, SP: specificity, AUC: area under the curve.

Following model training using binary cross-entropy with logits loss, EditFinder achieves an AUC of 0.9970 ± 0.0004 on the development set and 0.9813 ± 0.0021 on the independent test set for 111-nt sequences (Figure 1C–D). DeepLIFT [11] analysis further reveals sequence features contributing to the model predictions, including a preference for guanine (G) at the 3′ position immediately downstream of the editing site (Figure 1E), consistent with the known sequence preferences of ADARs [12]. These results demonstrate that EditFinder can accurately distinguish A-to-I RNA editing sites from genomic A-to-G SNPs based on their local sequence contexts.

To determine the optimal input sequence length and alignment weighting factor, an exhaustive evaluation was performed using ten-fold cross-validation on the development set (Figure 2A-B). The optimal configuration consists of an input sequence length of 111 nt and an alignment weighting factor (α) of 279. Using this configuration, the effects of sequence encoding and the contributions of the convolutional and alignment-based components were further evaluated. The 3-mer encoding achieves significantly higher predictive performance than the 1-mer encoding (Figure 2C). Ablation analyses further evaluated the contributions of the two major components of EditFinder. Models using only convolutional sequence features or only sequence alignment information show significantly lower performance than the full model integrating both feature types (Figure 2D–E). Together, these results support the selected configuration of EditFinder, including an input sequence length of 111 nt, an alignment weighting factor (α) of 279, 3-mer encoding, and the integration of convolutional and alignment-based sequence features.

**Figure 2.**
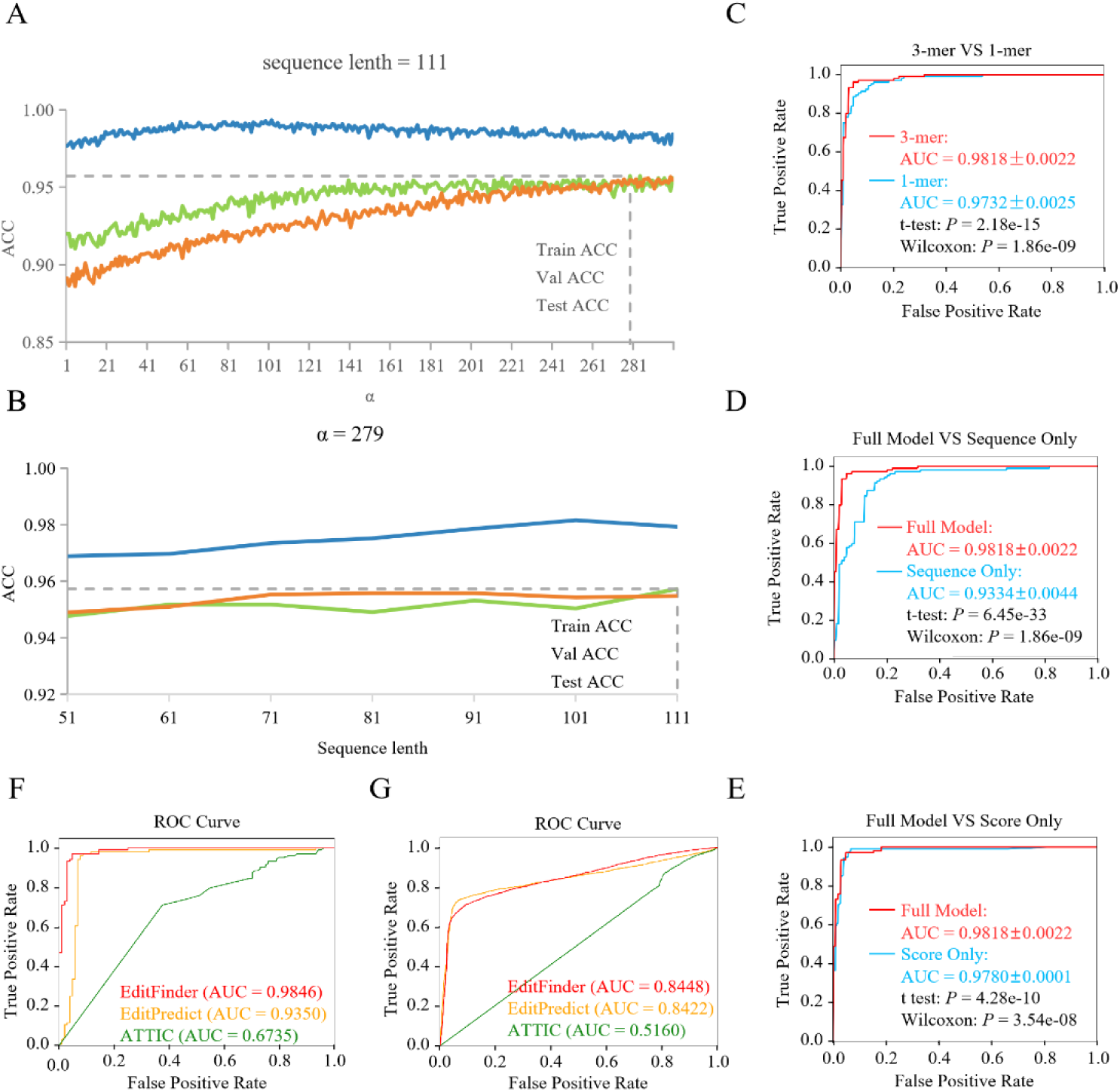
Evaluation of the EditFinder model. (A) Selection of the optimal alignment weighting factor (α) using a sequence length of 111 nt. (B) Selection of the optimal sequence length using an alignment weighting factor (α) of 279. (C) Comparison of EditFinder using 1-mer and 3-mer sequence encoding. (D) Ablation analysis comparing the sequence-only model with EditFinder. (E) Ablation analysis comparing the alignment score-only model with EditFinder. (F) Comparison of EditFinder with two published tools, EditPredict and ATTIC, using the test dataset constructed according to the pipeline shown in Figure 1. (G) Comparison of EditFinder with EditPredict and ATTIC using sequencing reads constructed according to the pipeline shown in Figure S1.

Moreover, EditFinder was compared with two existing A-to-I RNA editing prediction methods, EditPredict [13] and ATTIC [14], using the test set constructed for model evaluation (Figure 1A) and sequences reconstructed from RNA editing sites and SNPs identified in matched RNA-seq and DNA-seq data from the same sample (Figure S1). EditFinder shows significantly higher predictive performance than both EditPredict and ATTIC across the two datasets (Figure 2F-G). The improved performance may be attributed to two key features of EditFinder. First, the sequences used for model development are stringently filtered to directly model the key sequence-level differences between RNA editing and SNP-derived nucleotide changes. Second, EditFinder captures both local sequence patterns and sequence-level similarity for A-to-I RNA editing detection. These results demonstrate the superior performance of EditFinder.

### 2.2. Genome-wide distribution of A-to-I RNA editing sites identified by EditFinder

Given the high predictive performance of EditFinder in A-to-I RNA editing site detection, the model was applied to scan the whole-genome reference sequence. A total of 122,400,020 candidate RNA editing sites were identified, of which 66.74% had a predicted probability above 0.9 (Figure S2). The predicted sites are provided in GTF format through Synapse (syn78022039). Approximately half of the predicted sites are located within protein-coding genes, whereas 99.55% are located in non-coding regions (Figure 3A). Notably, 87.5% of the predicted sites are located within Alu elements (Figure 3B). This genomic distribution is consistent with previous studies reporting preferential localization of A-to-I RNA editing sites in non-coding regions and Alu elements [15], supporting the reliability of the predicted editing sites.

**Figure 3.**
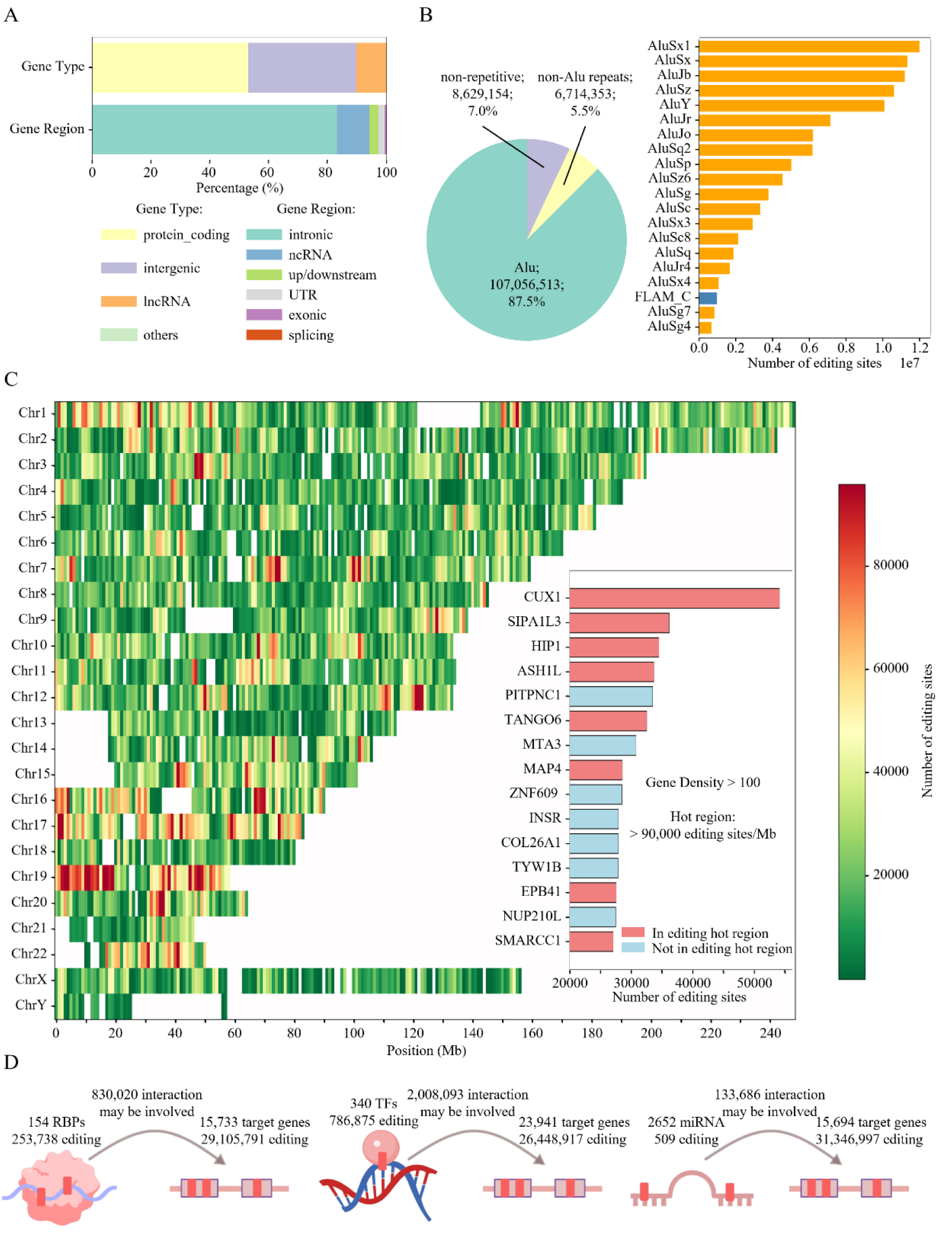
Genomic annotation of RNA editing sites. (A) Distribution of editing sites across gene type (upper) and gene regions (lower). (B) Overlap of RNA editing sites with repetitive elements. Left: Proportional distribution of RNA editing sites located within Alu elements, non-Alu repetitive elements, and non-repetitive genomic regions. Right: Top 20 repeat classes ranked by the number of overlapping RNA editing sites. (C) Chromosomal distribution and gene density features of editing events. Gene density is defined as the number of editing sites per 1kb of gene body length. (D) Functional regulatory annotation of RNA editing sites associated with RNA-binding proteins (RBPs), transcription factors (TFs) and microRNAs (miRNAs).

The density of predicted RNA editing sites was further examined across the genome. Regions with high editing site density are observed on chromosome 19 (Figure 3C), consistent with the reported high abundance of double-stranded RNA structures on this chromosome [16]. Genes with the highest editing-site densities include CUX1, SIPA1L3, HIP1, ASH1L, and PITPNC1 (Figure 3C). Notably, PITPNC1 was also reported as the gene with the highest enrichment of RNA editing sites in endometrioid ovarian cancer [8]. These genes represent diverse functional categories, including transcriptional and chromatin regulation [17; 18], cell polarity and cytoskeletal organization [19], and intracellular trafficking [20; 21]. Moreover, the predicted editing sites are widely distributed across RNA-binding proteins (RBPs), transcription factors (TFs), and microRNAs (miRNAs), as well as their respective target genes, based on annotations from RBP-Tar [22], the UCSC database [23], and TargetScan version 8.0 [24], respectively (Figure 3D). Editing at these sites may influence transcriptional regulatory networks by altering RNA sequence and structure or modulating interactions with regulatory factors. Dysregulated A-to-I RNA editing may consequently contribute to disease development and progression. Together, these results provide a genome-wide map of candidate A-to-I RNA editing sites and highlight their broad distribution across genomic regions and regulatory networks.

### 2.3 EditFinder uncovers a comprehensive landscape of SNP effects on A-to-I RNA editing

Previous studies have highlighted the potential SNP effect on RNA editing regulation [9; 25], suggesting that sequence variations may alter the recognition and editing propensity of RNA editing sites. Following the establishment of a genome-wide A-to-I RNA editing landscape, EditFinder was further applied to systematically assess the effects of individual SNPs on sequence-dependent RNA editing prediction. In total, 2,792,887,724 SNP-editing sequence pairs were analyzed. The effects of SNPs were assessed by comparing predicted RNA editing probabilities between reference and SNP-altered sequences. This analysis provides a genome-wide assessment of the potential impact of genetic variation on A-to-I RNA editing.

The magnitude of changes in predicted editing probability generally decreases with increasing distance between the SNP position and the RNA editing site (Figure 4A, syn78022039), suggesting that SNPs located closer to the editing site have a greater impact on editing probability. Among all examined positions, substitutions at the 3′-adjacent position exert the strongest effects, affecting the largest number of SNP-editing sequence pairs (Figure 4B). Specifically, substitutions of the native G or C nucleotides at this position with other nucleotides predominantly decrease predicted editing probability, whereas substitutions of the native T nucleotide with others predominantly increase it (Figure 4C). Notably, substitutions between G and C (G-to-C or C-to-G) or between A and T (A-to-T or T-to-A) result in smaller changes in editing probability than substitutions between G/C and A/T (Figure 4C). These observations are consistent with the reported ADAR nucleotide preference at the nearest downstream position (G > C > A > U) [12]. At the 5′-adjacent position, substitutions of the native G nucleotide with others predominantly increase predicted editing probability (Figure 4D), consistent with the reported lower ADAR preference for G at this position [12]. Furthermore, SNPs that disrupt preferred trinucleotide contexts, characterized by non-G nucleotides at the upstream position and G/C at the downstream position, generally lead to substantial reductions in predicted editing probability (Figure 4E). Among the 15 SNP-associated triplet changes showing the largest effects on editing probability, 12 (80%) involve substitutions at the downstream position (Figure 4F). Together, these results reveal the strong position- and nucleotide-dependent effects of SNPs on A-to-I RNA editing prediction, highlighting the prominent contribution of the 3′-adjacent nucleotide to sequence-dependent A-to-I RNA editing prediction.

**Figure 4.**
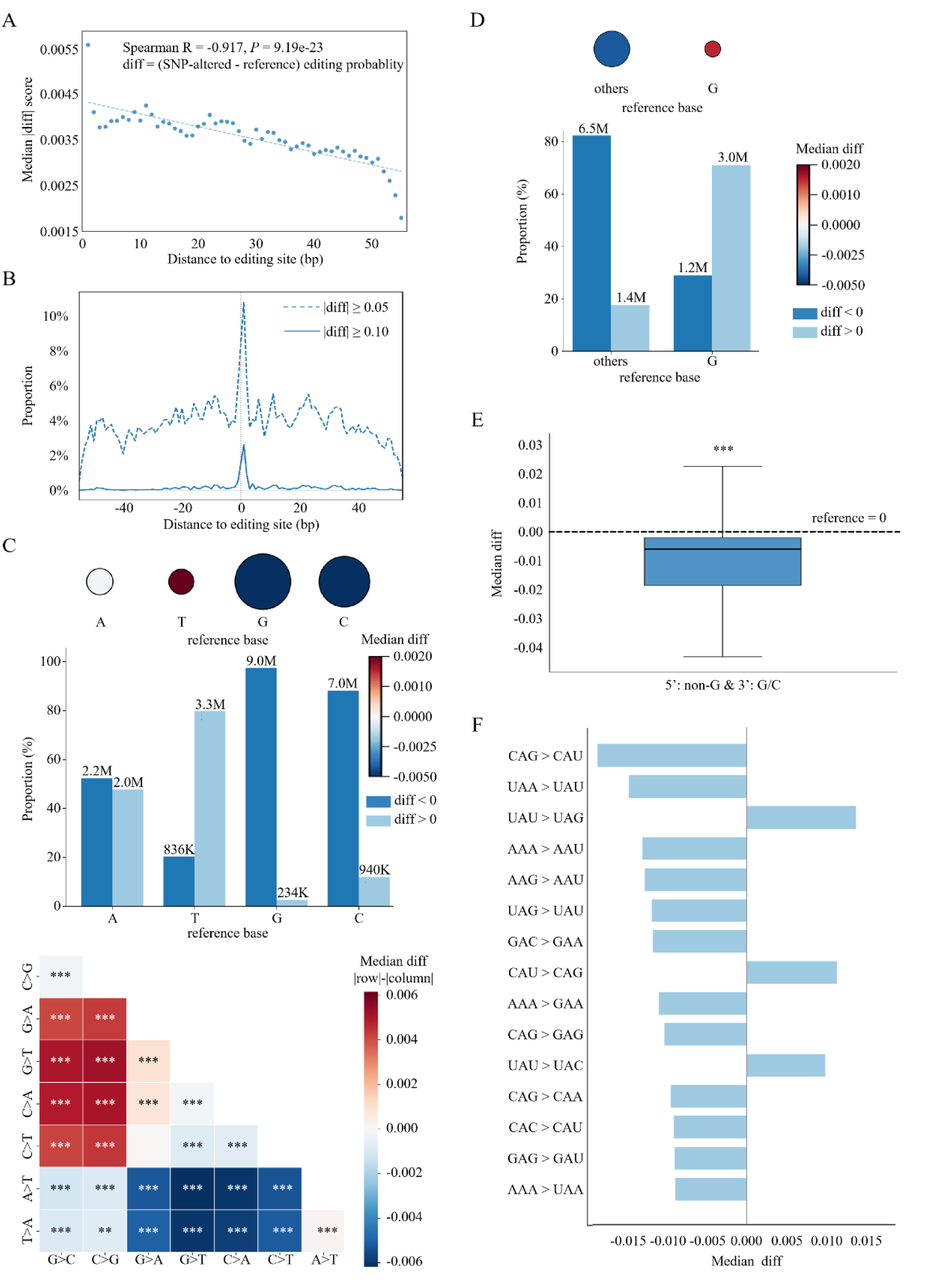
Genome-wide characterization of SNP effects on A-to-I RNA editing prediction. (A) Relationship between the distance of SNP positions from the RNA editing site and the median absolute difference in predicted editing probability between SNP-altered and reference sequences. (B) Proportions of SNP-editing sequence pairs showing changes above the indicated absolute difference thresholds across different SNP positions relative to the editing site. (C) Effects of nucleotide substitutions at the 3’-adjacent position of the editing site. Upper panel: distribution of reference nucleotides with bubble size representing the number of pairs and color indicating the median difference in predicted editing probability between SNP-altered and reference sequences. Middle panel: proportions of sequence pairs showing decreased or increased predicted editing probability for each reference nucleotide. Lower panel: comparison of changes in predicted editing probability associated with individual nucleotide substitutions. (D) Effects of nucleotide substitutions at the 5’-adjacent position of editing site. Upper panel: distribution of reference nucleotides. Lower panel: proportions of sequence pairs showing decreased or increased predicted editing probability for reference nucleotide. (E) Effects of SNPs on preferred trinucleotide sequence contexts, characterized by non-G nucleotides at the 5’ position and G/C at the 3’ position. (F) 15 trinucleotide substitutions associated with the largest changes in median predicted editing probability. ***: P<0.001, **: P<0.01.

### 2.4 SNPs alter the predicted RNA editing potential of sites in the REDIportal resource

RNA-seq-based RNA editing detection often relies on established reference resources, such as REDIportal [7], to identify known A-to-I RNA editing sites. However, these resources may not fully account for genomic sequence variations in individual samples. Because SNPs can alter the local sequence context surrounding an editing site [8], they may affect the reliability of RNA editing predictions when reference resources are applied to samples with different genomic backgrounds. To investigate this effect, EditFinder was used to evaluate sites in REDIportal based on both reference genome sequences and sequences carrying SNPs.

Using reference genome sequences, EditFinder classified 90.31% of the evaluated REDIportal sites as RNA editing sites (Figure 5A). Among the remaining sites, almost all had at least one SNP annotated within their surrounding sequences (Figure 5B). To further assess the potential effects of these variants, individual SNPs were introduced into the corresponding sequences, and the resulting sequences were reevaluated. Among the sites classified as SNPs by EditFinder using reference sequences, 99.87% showed an increased predicted RNA editing probability following the introduction of the best single SNP (Figure 5C). The distribution of these best SNPs across the surrounding sequence regions is shown in Figure 5D. Notably, SNPs at 3’-adjacent position of the candidate editing site are particularly likely to increase the predicted RNA editing probability, consistent with the known sequence preferences of ADARs [12].

**Figure 5.**
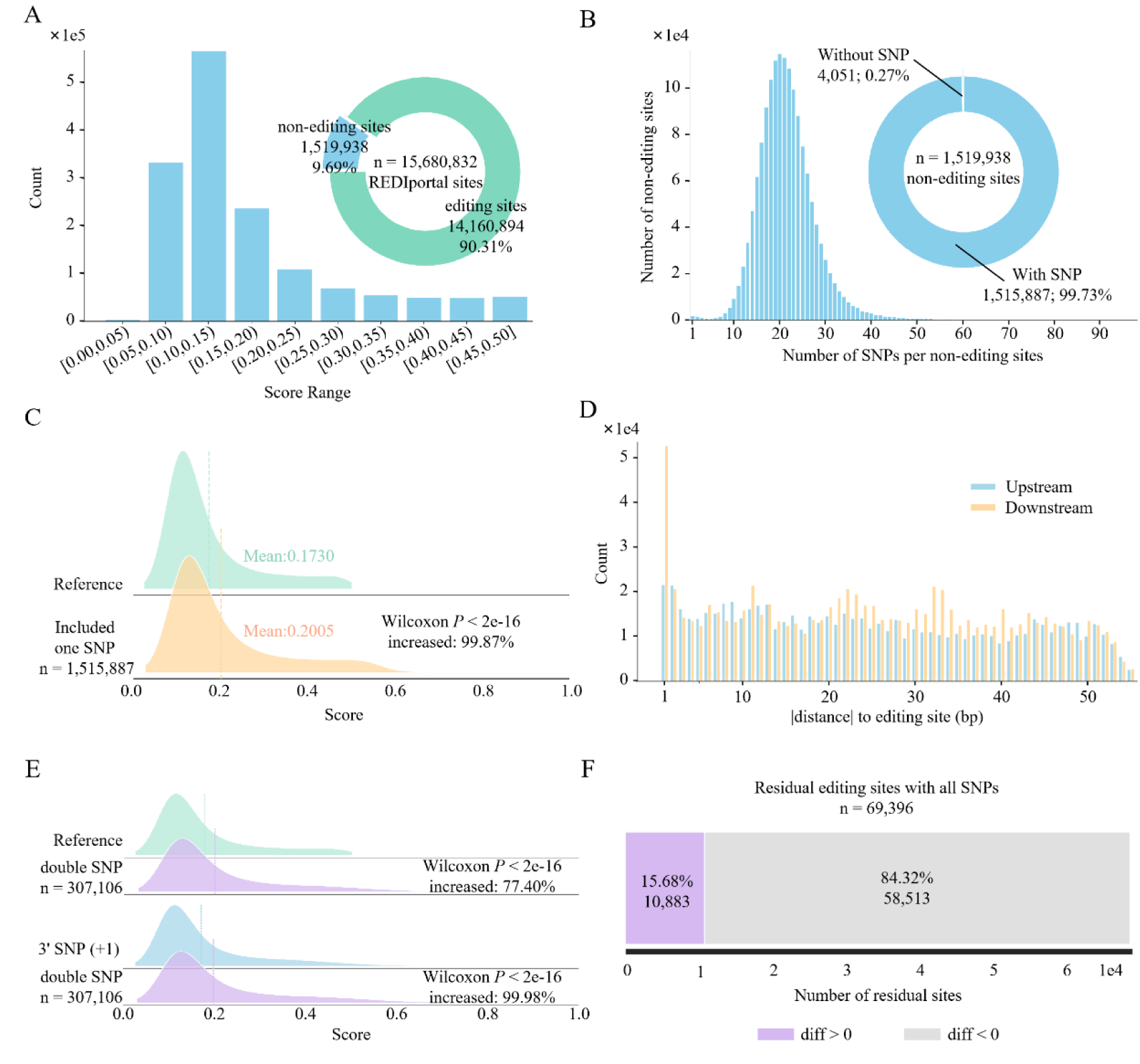
Effects of SNPs on the predicted RNA editing potential of sites in REDIportal. (A) Distribution of EditFinder prediction scores for REDIportal sites using reference genome sequences. (B) Distribution of the number of SNPs in the surrounding sequences of the sites classified as non-editing based on reference sequences. (C) Comparison of EditFinder prediction scores between reference sequences and sequences incorporating the single SNP associated with the highest prediction score for each site. (D) Positional distribution of SNPs shown in panel C within the flanking regions of RNA editing sites. (E) Comparison of prediction scores between reference sequences and sequences carrying an additional SNP at a position other than 3’-adjacent position of the editing site. (F) Distribution of prediction score differences for sites that did not show an increased score in panel E following the introduction of all SNPs.

The potential effects of additional SNPs were further examined in sites carrying a SNP at the 3’-adjacent position, which increased the predicted RNA editing probability relative to the reference sequence in 45.82% of cases. Following the introduction of an additional selected SNP into the surrounding sequence, 77.40% of these sites showed higher predicted RNA editing probabilities than the reference sequence (Figure 5E). Among the remaining sites, 15.68% sites also exhibited increased probabilities when additional SNPs were introduced. These findings indicate that SNPs in the local sequence context can substantially alter EditFinder predictions, particularly when variants modify nucleotides at positions associated with ADAR sequence preferences. Therefore, RNA editing events identified solely through reference database matching should be interpreted with caution, particularly when disease-specific genomic variation occurs around the predicted RNA editing sites.

### 2.5 EditFinder captures trends in SNP-associated RNA editing frequency changes

The preceding analyses demonstrate the utility of EditFinder in identifying candidate A-to-I RNA editing sites and assessing the effects of SNPs on their predicted editing potential. Notably, many SNPs induce relatively subtle changes in predicted editing probabilities without resulting in the gain or loss of predicted editing sites, raising the possibility that these changes may reflect alterations in RNA editing frequency. To investigate this possibility, SNP-associated changes in EditFinder prediction scores were integrated with the corresponding cis-edQTL effect sizes retrieved from GPEdit [25]. EditFinder-derived prediction changes are significantly associated with edQTL effect sizes (Figure 6A). Furthermore, SNP–editing sequence pairs with increased predicted editing probabilities are associated with positive edQTL effect sizes, and the magnitude of these changes is significantly greater than that observed for pairs with decreased predicted editing probabilities (Figure 6B).

**Figure 6.**
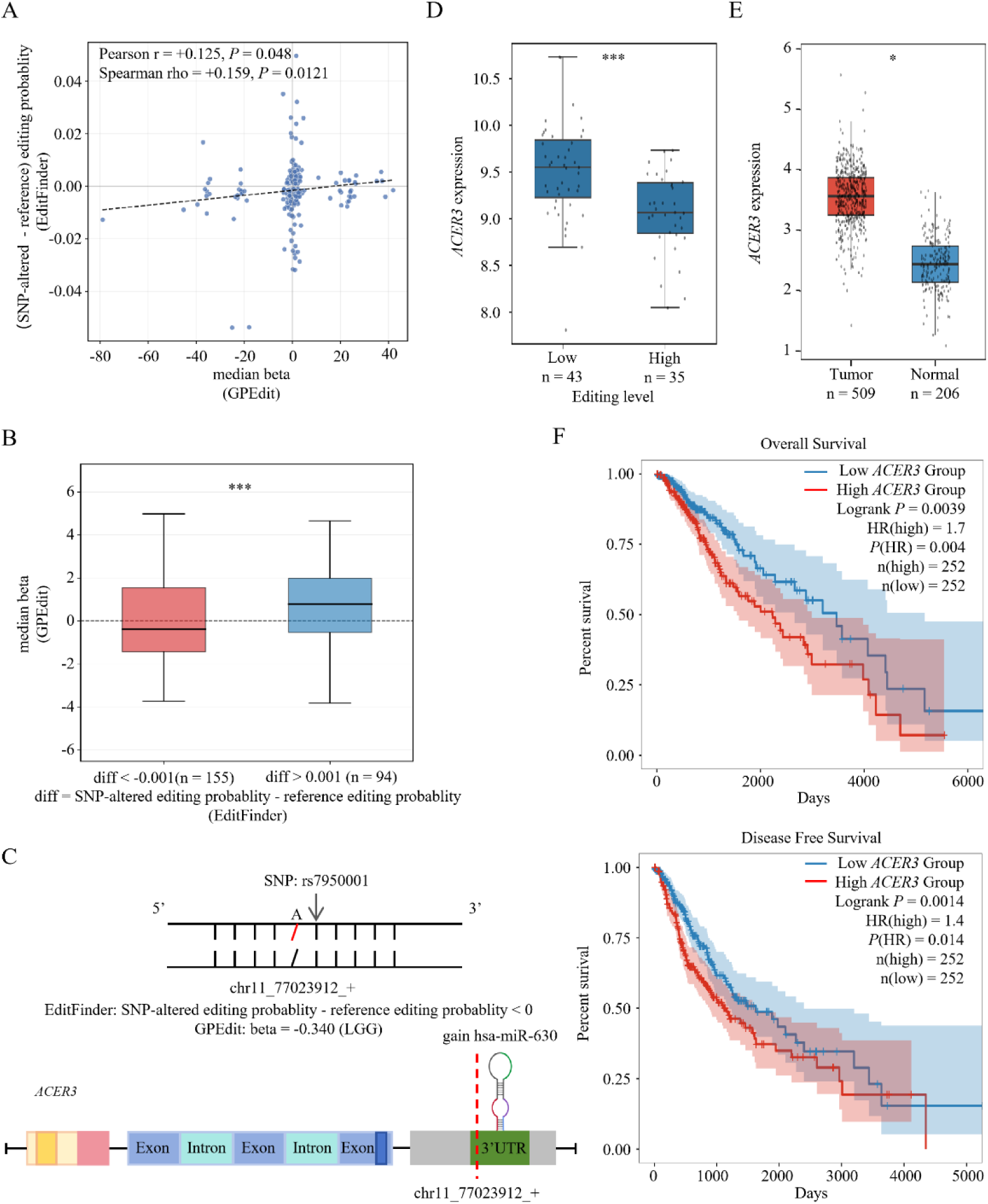
EditFinder captures trends in SNP-associated RNA editing frequency changes and identifies potential disease-associated regulatory mechanisms. (A) Correlation between SNP-associated changes in EditFinder-predicted RNA editing probabilities and median effect sizes (beta) estimated by editing quantitative trait locus analysis across cancer types from the GPEdit database. (B) Comparison of beta values between SNP-editing sequence pairs with increased and decreased RNA editing probability changes. (C) Schematic illustration of a 3’-adjacent SNP in *ACER3* associated with reduced RNA editing frequency, which is predicted to result in decreased miRNA binding (CAeditome). (D) Comparison of *ACER3* expression between samples with high and low levels of RNA editing. (E) Comparison of *ACER3* expression between tumor and normal samples. (F) Kaplan-Meier survival analyses showing the associations between *ACER3* expression and overall survival or disease-free survival.

Further analysis of a SNP located at the 3’-adjacent position of an RNA editing site in *ACER3* reveals a potential regulatory link between genetic variation, RNA editing, and gene expression in cancer (Figure 6C) [26]. This SNP is associated with a reduction in RNA editing prediction probability (Figure 6C), while the corresponding editing event is predicted to mediate miRNA regulation that may suppress *ACER3* expression (Figure 6C-D, Figure S3). Given the higher expression of *ACER3* in tumor samples than in normal controls (Figure 6E), together with the association between elevated *ACER3* expression and poorer overall and disease-free survival (Figure 6F), this SNP may represent a disease-associated variant with a potential RNA editing-mediated regulatory effect. Collectively, these findings suggest that EditFinder-derived changes in editing probability can capture trends in SNP-associated RNA editing frequency and facilitate the identification of potential RNA editing-mediated regulatory mechanisms with disease relevance.

## 3. Discussion

This study presents EditFinder, a sequence-based model that demonstrates superior performance in A-to-I RNA editing identification compared with existing methods. This model provides a genome-wide resource of candidate A-to-I RNA editing sites based on the human reference genome, accompanied by comprehensive functional annotations at the gene, genomic region, repetitive element, and regulatory relationship. Furthermore, it enables a systematic genome-wide assessment of individual SNP effects on A-to-I RNA editing prediction, highlighting the prominent contribution of the 3’-adjacent nucleotide to sequence-dependent RNA editing. Additionally, its application to the REDIportal resource further underscores the importance of considering genetic variations when matching RNA-seq-derived candidate sites against catalogued RNA editing sites and interpreting RNA editing events. Finally, integrating SNP-associated changes in predicted editing probability, as assessed by EditFinder, with RNA editing quantitative trait associations suggests that EditFinder predictions may capture trends in SNP-associated RNA editing frequency variation. Collectively, this study establishes EditFinder as a framework for genome-wide A-to-I RNA editing identification and for exploring the potential influence of genetic variation on RNA editing.

The EditFinder-derived resource provides a comprehensive genome-wide reference of candidate A-to-I RNA editing sites. Compared with the existing REDIportal database, EditFinder identifies an additional 110,415,612 candidate editing sites, 64.9% of which have prediction probabilities above 0.9 (Figure S4). These sites are distributed across 32,721 genes, including 1,090 cancer-associated genes [27], suggesting the potential involvement of these editing sites in diverse biological processes and cancer-related mechanisms. The analysis highlights the value of this resource for investigating the functional landscape of RNA editing and its potential roles in diseases.

Moreover, EditFinder maps the effects of individual SNPs on A-to-I RNA editing prediction at the genome-wide scale (Figure 4) and assess SNP effects on REDIportal-documented editing sites whose reference sequences are predicted to be non-editing (Figure 5C-D). The results reveal that individual SNPs can affect the predicted editing probability of a candidate site by up to 0.9, potentially resulting in the loss or gain of RNA editing sites. Beyond individual SNP effects, the combined effects of selected double and multiple SNPs on RNA editing prediction were also investigated (Figure 5E-F). However, given the vast number of SNPs and the potentially enormous number of their combinations, exhaustive evaluation is computationally impractical. EditFinder therefore enables researchers to evaluate the effects of specific SNPs or SNP combinations on RNA editing at selected sites of interest.

Furthermore, although the association between EditFinder predictions and RNA editing quantitative trait associations suggests that sequence-based models may capture trends in RNA editing frequency variation, the current framework primarily focuses on local sequence features associated with editing probability. Future research may therefore focus on a more comprehensive characterization of RNA editing frequency variation by considering broader regulatory sequence contexts within specific disease settings. Such efforts may facilitate the identification of editing regulatory factors, provide further insights into the RNA editing-mediated mechanisms underlying disease development, and enable a more comprehensive assessment of SNP effects on disease-specific RNA editing frequency.

## 4. Material and Methods

### 4.1. Datasets involved in this study

To develop the A-to-I RNA editing identification model, RNA editing sites and genomic SNPs were collected from CAeditome [15] and dbSNP (Build 151), respectively. Their corresponding reference sequences were obtained from GENCODE (Version 32). Model validation using sequencing data was performed with paired RNA-seq (SRR21068783) and DNA-seq (SRR21068914) datasets, from which reads spanning candidate RNA editing sites and genomic variants were extracted. RNA editing sites catalogued in REDIportal [7] served as an external reference for evaluating the effects of SNPs on existing RNA editing resources. Significant SNP-editing site associations from GPEdit [25] were used to examine the relationship between SNP-associated changes in predicted editing probability and RNA editing quantitative trait effects. Finally, gene expression profiles and corresponding clinical information from the TCGA and GTEx projects were used to investigate the potential disease relevance of RNA editing-mediated regulatory mechanisms.

### 4.2. Construction of development and test datasets for A-to-I RNA editing detection model

To establish a sequence-based model for distinguishing A-to-I RNA editing from SNP-derived nucleotide changes, reference genome sequences corresponding to reliable RNA editing sites and genomic A-to-G SNPs were collected and subjected to stringent filtering to minimize potential ambiguity between the two classes (Figure 1A). A-to-I RNA editing sites obtained from the CAeditome database were used to construct the positive set. Editing sites that overlapped with SNPs annotated in dbSNP (version 151) or had more than 10 missing values were removed. The remaining editing sites were further filtered to exclude sites whose surrounding upstream or downstream regions contained SNPs. Reference genome sequences centered on the remaining editing sites were then extracted for model development. Sequence redundancy was controlled by removing sequences with a pairwise sequence similarity greater than 0.95. A-to-G SNPs obtained from dbSNP were used to construct the negative set. SNPs that overlapped with known RNA editing sites or whose surrounding upstream or downstream regions contained additional SNPs were excluded. Reference genome sequences centered on the remaining SNPs were then extracted, and an equal number of negative sequences were randomly selected to match the number of positive sequences.

To assess model performance on independent genomic regions, RNA editing and SNP sequences located on chromosomes 1 and 19 were reserved as the independent test set, whereas sequences from the remaining chromosomes were used for model development. The test set was not used during model development or parameter optimization. The development dataset was subjected to ten-fold cross-validation, with the data in each fold further divided into training and validation sets for hyperparameter optimization. Model performance was evaluated using accuracy (ACC), sensitivity (SE), specificity (SP), and the area under the receiver operating characteristic curve (AUC).

### 4.3. Model architecture

The input sequences were encoded using a 3-mer representation, resulting in an input feature matrix of size 4^3^×(*L*-2) where *L* denotes the sequence length. The encoded sequences were then processed through two convolutional layers with ReLU activation and dropout regularization. The first and second convolutional layers generated 128 and 256 feature maps, respectively.

In parallel with convolutional feature extraction, each input sequence was aligned against all positive sequences in the development set to calculate sequence alignment scores using Needleman-Wunsch method [28]. The maximum alignment score was selected as an additional sequence similarity feature. For each positive sequence in the development set, its own sequence was excluded from the reference set when calculating the maximum alignment score to avoid self-alignment. The resulting maximum alignment score was multiplied by a weighting factor α and then integrated with the convolutional features. The resulting combined features were flattened and passed through two fully connected layers with 256 and 128 units, respectively. Finally, the output layer with sigmoid activation generated a predicted probability of A-to-I RNA editing ranging from 0 to 1, with higher values indicating a higher predicted probability of RNA editing.

### 4.4. Identification of RNA editing sites and SNPs from RNA-seq and DNA-seq data

To evaluate EditFinder using sequencing data from biological samples, RNA editing sites and genomic SNPs were identified from matched RNA-seq (SRR21068783) and DNA-seq datasets (SRR21068914) following the pipeline shown in Figure S1. For RNA-seq data, sequencing reads were aligned to the reference genome using combined HISAT2 and BWA [29], whereas DNA-seq reads were aligned to the same reference genome using BWA and subsequently pre-processed with the Genome Analysis Toolkit (GATK). Alignments with a mapping quality score below 10, more than six mismatches or a base quality score below 18 were excluded. The resulting BAM files were used for downstream variant and RNA editing analyses.

RNA editing sites and SNPs were identified using REDItools by jointly analyzing the RNA-seq and DNA-seq BAM files. Candidate sites were selected based on the following criteria: an RNA-seq read depth of at least 10, a mean base quality of at least 20, an editing frequency of at least 5%, at least three editing-supporting reads, and a DNA-seq read depth of at least 10. Sites not annotated in REDIportal were excluded to obtain a high-confidence set of known RNA editing sites. For genomic SNPs, only A-to-G substitutions were retained for subsequent analyses.

For each retained RNA editing site and SNP, the corresponding sequence context was reconstructed from the RNA-seq data. The RNA-seq BAM file was used to extract reads spanning the candidate sites, and Trinity was subsequently used to assemble the local sequences into contigs. Contigs were retained only when they covered the candidate site and provided sufficient sequence coverage. Contigs shorter than the optimal input length determined for EditFinder or with a k-mer coverage below 1 were excluded. The resulting contigs were aligned to the reference genome using HISAT2, and contigs that could not be mapped to the reference genome or had an alignment score below 2 were excluded. The sequences surrounding the retained candidate sites were then extracted from the mapped contigs to generate the final SNP and RNA editing sequence sets for evaluation of EditFinder.

### 4.5 Analysis of SNP effects on RNA editing site prediction

To investigate the effects of SNPs on RNA editing site prediction, reference sequences surrounding candidate editing sites were modified by introducing SNPs at individual positions, and the resulting sequences were evaluated using EditFinder. The change in predicted RNA editing probability relative to the original reference sequence was calculated. The positional distribution of SNPs associated with changes in predicted RNA editing probability was also examined to identify sequence positions with potentially greater effects on model predictions.

## 5. Data availability

RNA editing sites and SNPs were retrieved from CAeditome and dbSNP (Build 151) respectively. Reference genome sequences were obtained from GENCODE (Version 32). Paired RNA-seq and DNA-seq data were obtained from SRR21068783 and SRR21068914 respectively. The genome-wide reference A-to-I RNA editing resource predicted by EditFinder is available through Synapse (syn78022039). Human RNA editing site annotations from REDIportal were used as an external reference dataset. Statistical edQTL results were obtained from GPEdit. *ACER3* gene expression and associated clinical data were obtained from TCGA and GTEx project.

## 6. Code availability

All code associated with this study is available at https://github.com/swu13/EditFinder/.

## 7. Supplementary Data

Supplementary files are available online along with the manuscript.

## 8. Acknowledgement

This work was supported by the Fundamental Research Funds for the Central Universities, the National Natural Science Foundation of China (Grant No. 82227802, 62002270), Key Research and Development Program of Shaanxi Province (No. 2026SF-YBXM-279), and Science and Technology Development Incubation Fund Program of Shaanxi Provincial People’s Hospital (No.2023YJY-18).

