## Supplementary Figures1-4 and Supplementary Table 1 for "EditFinder: A Sequence-Based Model for A-to-I RNA Editing Identification and Characterization of Genetic Variant Effects"

Table S1. Sample sizes of the development and test sets for sequences of different lengths.

| | **51nt** | **61nt** | **71nt** | **81nt** | **91nt** | **101nt** | **111nt** |
|---|---|---|---|---|---|---|---|
| **Development Set** | 1,612 | 1,220 | 977 | 742 | 597 | 468 | 363 |
| **Test Set** | 440 | 343 | 272 | 201 | 163 | 129 | 104 |

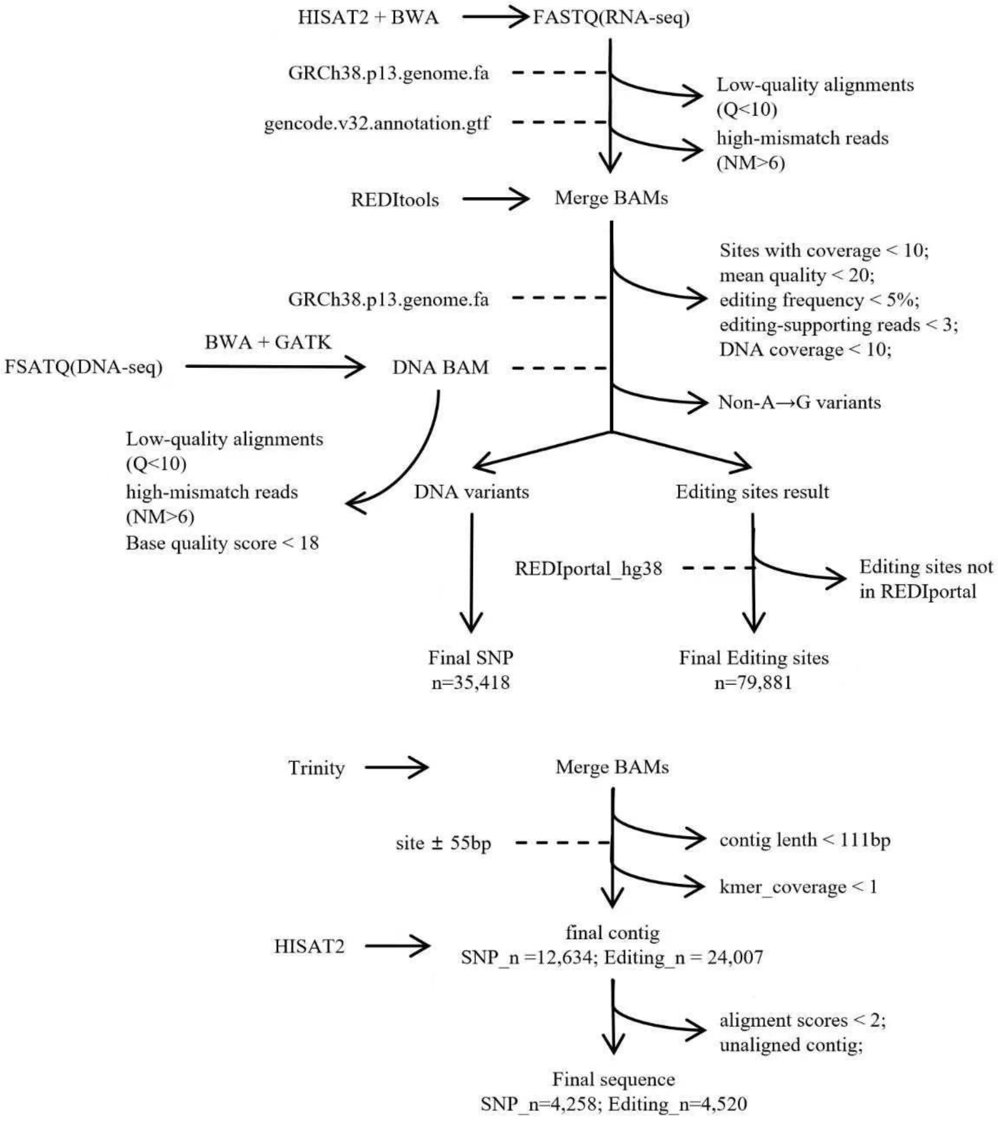


Figure S1. Extraction pipeline for sequencing reads from sample AUR-AFUG-TTM1, sequenced by both RNA-seq (SRR21068783) and DNA-seq (SRR21068914).

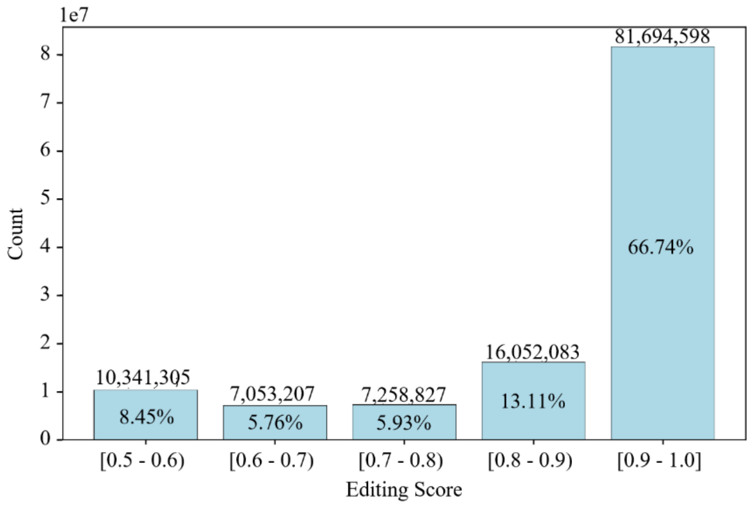


Figure S2. Distribution of predicted A-to-I RNA editing sites across prediction probability intervals.

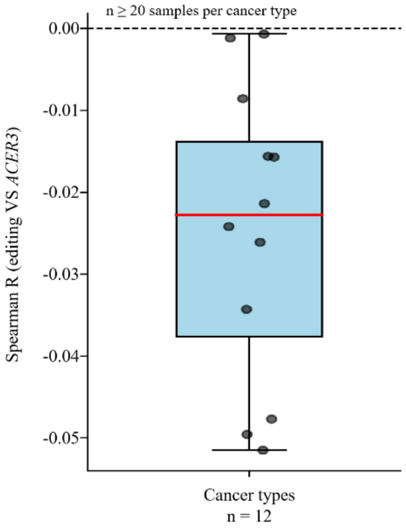


Figure S3. Boxplots showing Spearman correlation coefficients (R) between RNA editing levels and *ACER3* expression across different cancer types.

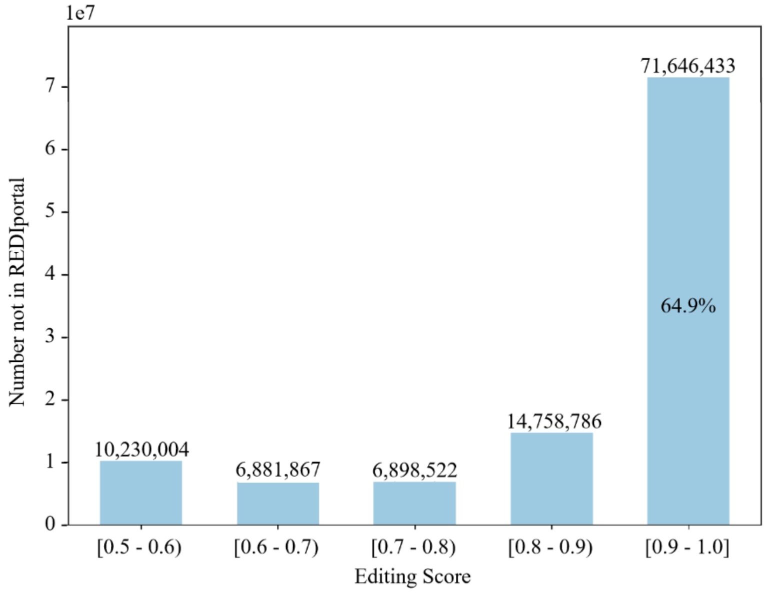


Figure S4. Distribution of predicted A-to-I RNA editing sites absent from the REDIportal database across prediction probability intervals.
